# Accelerating Genome- and Phenome-Wide Association Studies using GPUs – A case study using data from the Million Veteran Program

**DOI:** 10.1101/2024.05.17.594583

**Authors:** Alex Rodriguez, Youngdae Kim, Tarak Nath Nandi, Karl Keat, Rachit Kumar, Rohan Bhukar, Mitchell Conery, Molei Liu, John Hessington, Ketan Maheshwari, Drew Schmidt, VA Million Veteran Program, Edmon Begoli, Georgia Tourassi, Sumitra Muralidhar, Pradeep Natarajan, Benjamin F Voight, Kelly Cho, J Michael Gaziano, Scott M Damrauer, Katherine P Liao, Wei Zhou, Jennifer E Huffman, Anurag Verma, Ravi K Madduri

## Abstract

The expansion of biobanks has significantly propelled genomic discoveries yet the sheer scale of data within these repositories poses formidable computational hurdles, particularly in handling extensive matrix operations required by prevailing statistical frameworks. In this work, we introduce computational optimizations to the SAIGE (Scalable and Accurate Implementation of Generalized Mixed Model) algorithm, notably employing a GPU-based distributed computing approach to tackle these challenges. We applied these optimizations to conduct a large-scale genome-wide association study (GWAS) across 2,068 phenotypes derived from electronic health records of 635,969 diverse participants from the Veterans Affairs (VA) Million Veteran Program (MVP). Our strategies enabled scaling up the analysis to over 6,000 nodes on the Department of Energy (DOE) Oak Ridge Leadership Computing Facility (OLCF) Summit High-Performance Computer (HPC), resulting in a 20-fold acceleration compared to the baseline model. We also provide a Docker container with our optimizations that was successfully used on multiple cloud infrastructures on UK Biobank and All of Us datasets where we showed significant time and cost benefits over the baseline SAIGE model.

## Introduction

The rapid expansion of biobanks has significantly advanced genomic discoveries, facilitating studies on the genetic basis of disease and catalyzing studies in personalized medicine. The increasing number of newly formed biobanks, alongside the continued growth of established ones, enables researchers to conduct studies with increasingly larger sample sizes, yielding more robust and generalizable findings. Furthermore, biobanks linked to electronic health records (EHR) have been instrumental in translational studies, providing data on both the genome and phenome in large populations (1–7).

However, the unprecedented size of the data accessible via biobanks require researchers to consider the computational challenges arising from data complexity, analysis methodologies, statistical frameworks, and infrastructure constraints. Addressing the computational limitations requires development of innovative algorithms, optimization strategies, and adaptable computing architectures tailored to the unique requirements of biomedical research. Additionally, fostering collaboration among computational scientists, statisticians, and domain experts proves indispensable in crafting resilient computational tools and workflows capable of facilitating the efficient analysis of burgeoning biobank data. Leveraging the enhanced computational infrastructures afforded by high-performance computing (HPC) and cloud environments further augments the capacity for comprehensive analysis within the biomedical domain.

One routine statistical analysis that utilizes genetic and phenotypic data available from large biobanks is the genome-wide association study (GWAS). The purpose of GWAS is to identify association between polymorphic DNA variants in the genome among biobank participants and a phenotypic trait or disease of interest, typically extracted from EHR or clinical data from participants where the underlying computation involves millions of iterations of a generalized linear model over the available genetic exposures (8). Furthermore, the complexity of the analysis increases when state-of-the-art approaches extend the analysis to use multi-level models to better account for population architecture and relatedness, in addition to the volume of the data resulting in huge, dense matrix-matrix and matrix-vector operations that are performed as a part of statistical scaffolding (9–12). Compounding this computational complexity are data scaling challenges, including the desire to analyze the entire catalog of traits extracted from EHR data (i.e., the “phenome”, 100s-1000s of traits) and doing so across all population groups represented in order to capture the diverse representation that is increasingly available. The scale of this undertaking requires large amounts of disk space, fast processors, and innovative techniques to take advantage of the resources available.

The U.S. Department of Veterans Affairs (VA) Million Veteran Program (MVP) stands as a pioneering research endeavor, continually expanding in scope and offering researchers a platform to tackle the aforementioned challenges. This biobank, aimed at advancing precision medicine and improving healthcare outcomes for Veterans, includes a large number of individuals from underrepresented populations. The VA and the Department of Energy (DOE) established an Interagency Agreement (IAA) to combine VA’s vast array of clinical and genomic data with DOE’s national computing capabilities, including the most powerful supercomputer in the Nation, to push the frontiers of precision medicine and computing with vision to improve the lives of Veterans and all Americans and support the national Precision Medicine Initiative. Its primary mission revolves around furnishing comprehensive genotype-phenotype insights into prevalent and significant health outcomes, leveraging the most extensive EHR-linked biobank in the United States. Among the formidable computational challenges encountered in this research is the task of conducting GWAS across a staggering 3.5 billion genetic variants, spanning thousands of traits gleaned from the electronic health records of 635,969 MVP participants.

SAIGE (the Scalable and Accurate Implementation of Generalized Mixed Model algorithm) (10) is one such state-of-the-art multi-level modeling based GWAS approach designed to accommodate sample relatedness and manage unbalanced case-control ratios typical in biobanks like MVP. A crucial aspect of GWAS analysis entails constructing a Genetic Relationship Matrix (GRM), which measures the genetic relatedness or similarity among individuals within the study cohort. SAIGE offers users the option to generate either a sparse or a full GRM. While a sparse GRM boasts faster executions and lower memory demands, opting for a full GRM provides a more precise assessment of pairwise relatedness among all individuals, enhancing the depiction of genetic relationships. This proves invaluable for various downstream analyses, including estimating heritability, evaluating genetic correlations, and achieving a deeper understanding of the genetic architecture of the trait in question (14). To mitigate the computational burden associated with storing and inverting the full GRM, SAIGE employs the pre-conditioned conjugate gradient (15) method to iteratively solve linear equations. Nevertheless, it still faces substantial computation challenges due to extensive matrix and vector multiplications across numerous iterations.

In high-performance computing, processors face performance bottlenecks due to memory and disk input/output (I/O) operations. While processors execute computations rapidly, they depend on memory for data storage. Limited memory capacity necessitates frequent reads and writes to disk storage, slowing overall execution. Parallel processing distributes workloads across multiple compute processor units (CPU) but memory limitations persist. These bottlenecks are especially pronounced in large matrix operations. Graphics Processing Units (GPUs) provide an optimized architecture through high memory bandwidth and capacity (16), massive parallelism (17), and reduced data transfer between CPU and memory (18), particularly for large matrix operations. By leveraging these GPU attributes, computationally intensive tasks, like large matrix operations, can experience significance performance enhancements compared to CPU processing alone, mitigating memory and disk I/O bottlenecks.

We adapted SAIGE, initially tailored for CPU infrastructure, to utilize both CPUs and GPUs at the DOE Oak Ridge Leadership Computing Facility (OLCF) Summit HPC. This adaptation markedly expedited the analysis process, resulting in a more than 20-fold acceleration, far surpassing what would have been attainable on a CPU-based cluster. Furthermore, our work provided a *generic optimization framework* for other analytical tools based on generalized linear mixed models using a full GRM. We also created a Docker container for deployment on various cloud infrastructures. We present a comparison of the time and cost between the SAIGE GPU and CPU versions.

## Materials and Methods

### Study Design, Population Groups, and Phenotypic Definitions

The analysis involved a series of GWAS across 2,068 traits, covering a deep catalog of phenotypes extracted from EHR-derived diagnosis codes, clinical laboratory tests, vital signs, and survey responses. As previously described (13, 19, 20, 21, 22), the analysis was performed using data from 635,969 participants from MVP Genomics Release 4 (23) (**Table 1**) classified into four population groups based on genetic similarity (GIA) to the 1000 Genomes Project (24, 25) African (AFR, n = 121,177), Admixed Americans (AMR, n = 59,048), East Asian (EAS, n = 6,702), and European (EUR, n = 449,042) superpopulations. After imputation and quality control (QC) filtering, > 44.3M variants (minor allele count (MAC) ≥ 40) were included for analysis. For a visual representation of the analysis, please refer to **Figure 1a**, which illustrates the different quantities for which the analysis was conducted. After trait quality control, 1,854 binary and 214 quantitative traits were included in the downstream analysis in at least one population group (**Figure 1b**).

**Fig. 1.**
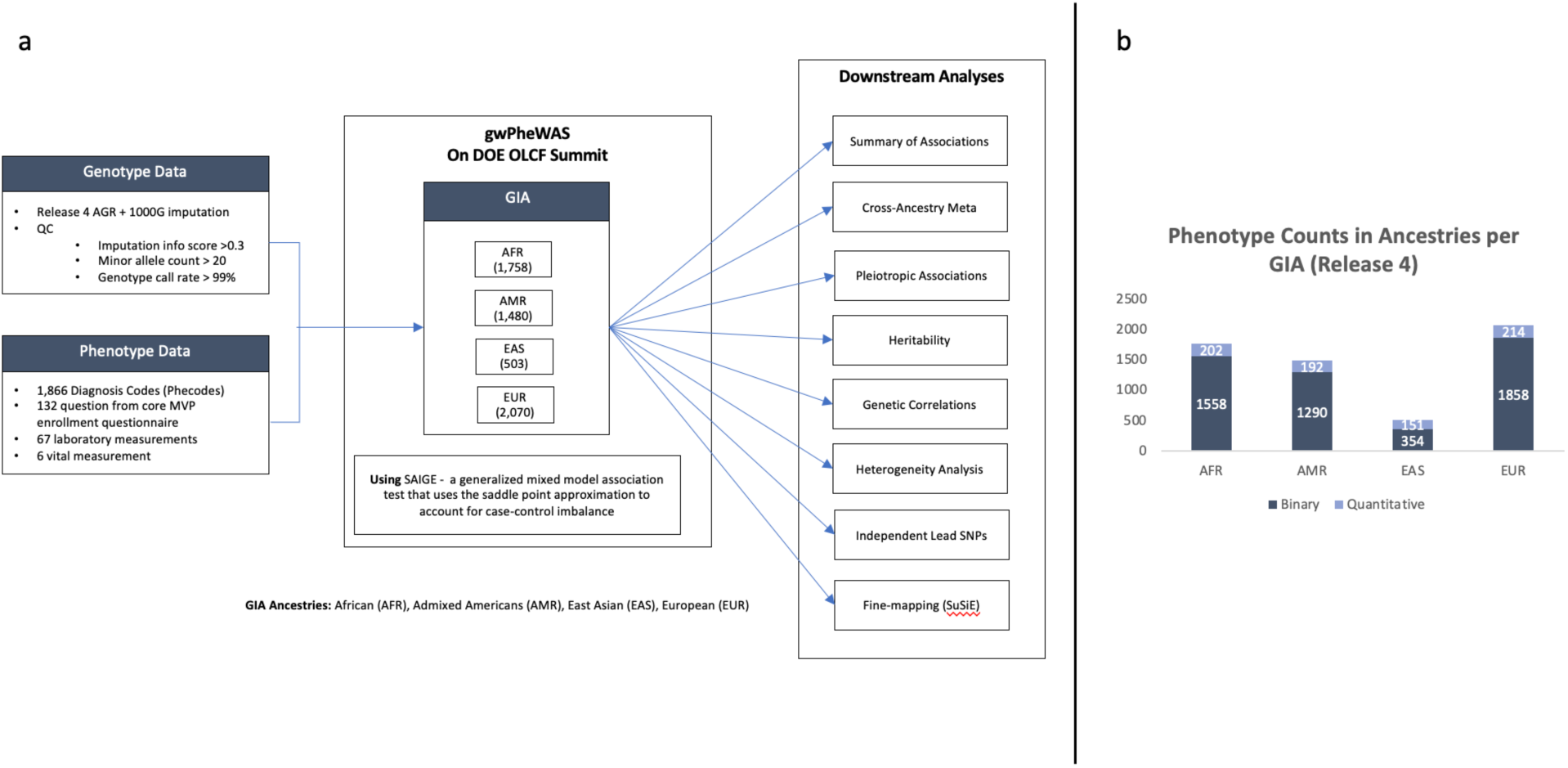
Overview of genomic analysis in multiple population groups. a) Schematic representation illustrating the diverse set of GIA population groups. The analysis covers a deep catalog of traits extracted from electronic health records, clinical laboratory tests, vital signs, and survey responses. b) Chart categorizing traits into binary or quantitative types across different population groups. The height of each bar corresponds to the number of traits in each category, providing an overview of the trait composition for subsequent genomic analyses.

**Table 1.**
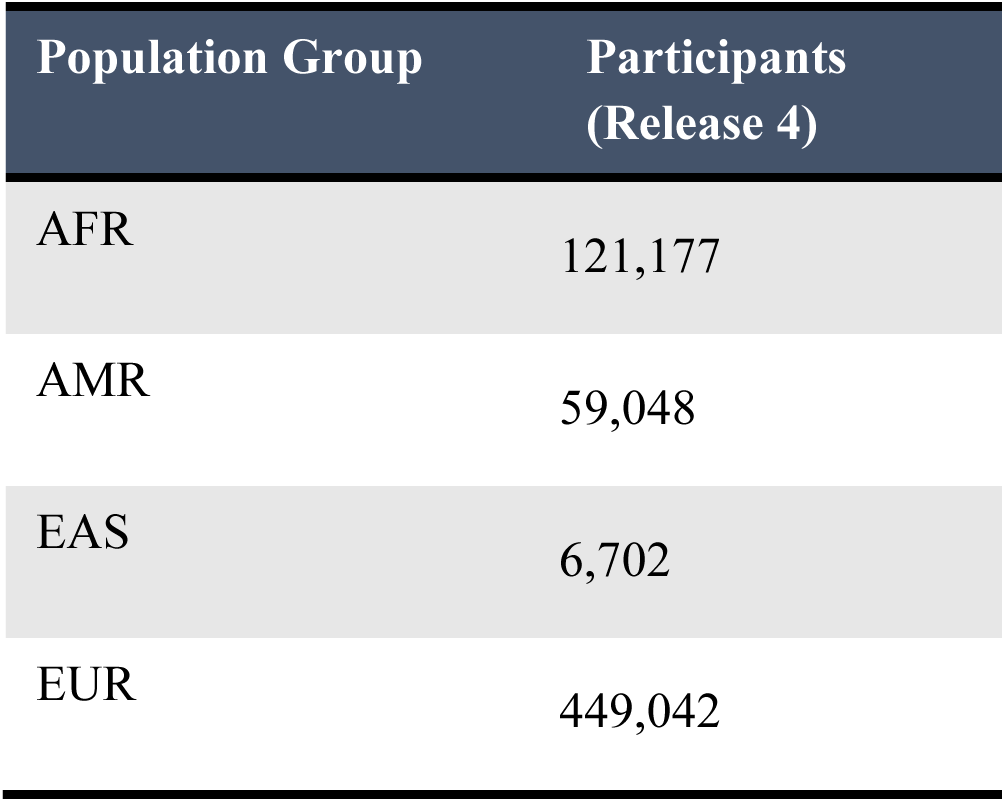
Participant quantity in each grouping method per population group. Data was made available on OLCF Summit HPC to perform a GWAS analysis for all traits analysis and all population groups.

Additionally, genotype data from the UK Biobank (3) and the All of Us Research Program (7) were utilized to test the software capabilities on a cloud environment. Within the UK Biobank data, we employed the European (EUR, n = 420,500) and African (AFR, n = 6,600) cohorts where PCA was used to measure the population structure (26). The All of Us cohorts included European (EUR, n = 133,000) and African (AFR, n = 55,000) population groups where PCA was used to measure the population structure (7).

### Biobank-scale genomic analysis across population groups

In total, 4,045 GWAS SAIGE runs were needed for the GW-PheWAS analysis which resulted in over 350 billion variant-trait associations across population groups. The current implementation of the SAIGE algorithm was not analytically tractable at this scale of computation. SAIGE uses R/C++ based tools developed for CPU environments and uses the Intel Threading Building Blocks (TBB) (27) library to enable parallelization. The SAIGE method comprises two primary steps. Step one involves fitting a linear/logistic mixed model with a GRM included under the null hypothesis that no genetic variants are associated with the phenotype of interest. We note that fitting the null mixed model involves thousands of matrix-matrix and matrix-vector operations, which are best suited for a GPU environment. Step two tests each SNP at a time across the genome for their association with the phenotype with a score test using the saddlepoint approximation (SPA) (28) and Firth regression methods (29) to account for unbalanced case-control sample sizes.

Directly genotyped variants were used for step one of SAIGE and were filtered for pairwise correlation with a window size, number of SNPs and VIF threshold of 50, 5, and 0.2 respectively using Plink1.9 (9). Imputed genetic dosages were used for step two of SAIGE. Only variants with an imputation quality > 0.3 and MAC ≥ 40 within the relevant population groups were included in the GWAS execution. Analyses were adjusted for age, sex, and the top ten population specific genetic principal components (PC) estimated by Principal component analysis (13).

### Computational Infrastructures

All GWAS analysis was conducted on the Summit HPC, located at DOE’s OLCF. It consists of 4,608 nodes, with each node featuring two IBM POWER9 processors and six NVIDIA Tesla V100 GPUs of 16 GB memory. All but 54 of the Summit nodes are equipped with 512 GB of DDR4 memory for the POWER9 processors and 96 GB of high-bandwidth memory (HBM2) for the V100 GPUs. The remaining 54 nodes in Summit HPC are high-memory nodes equipped with 2 TB of DDR4 memory for the CPUs and 192 GB of HBM2 for the GPUs with 32 GB of memory per GPU. Specifically, for the GW-PheWAS analysis, we utilized the nodes on Summit with 512 GB DDR4 memory and 96 GB HBM2 memory.

To further advance the use of SAIGE-GPU in various research environments, we generated Docker and Singularity containers. We conducted extensive testing of this containerized solution on the GPUs available on Google Cloud Platform (GCP) and Azure Cloud Platforms. Our code for containers and optimizations is provided (**Data and materials availability**).

## Results

We initially focused on optimizing step one for SAIGE as it can largely benefit by using GPUs for matrix-vector operations in the calculation of the GRM on the fly and employed MPI to distribute the data across multiple GPUs. While the standard SAIGE method on CPU-based machines was suitable for relatively small cohorts, it became impractical for larger population groups (e.g., groups similar to 1KG-European and 1KG-Africa in MVP) due to the substantial size of the matrix-vector operations involved. Employing the GPU-modified SAIGE framework, we successfully conducted a total of 4,045 independent GWAS runs. The GWAS analysis was accomplished within 14,286 GPU hours for step one, equivalent to 5 days of wall time, resulting in a 160-fold reduction in required core CPU hours in a CPU environment cluster (**Table 2**). Step two in SAIGE presented distinct challenges due to the need for millions of association tests for each trait, totaling over 3 billion association tests.

**Table 2.** Projection times to complete GWAS for all traits (5,819) using SAIGE step one using the different implementations of SAIGE: Native, OpenMP and GPU versions on CPU and GPU environments.

| Population Group | Trait Quantity | Step One CPU hours for all traits (Projected) |  | Step One GPU hours for all traits (Production) |
| --- | --- | --- | --- | --- |
|  |  | Native SAIGE | SAIGE-OpenMP | SAIGE-GPU |
| AFR | 1,760 | 322,768 | 78,266 | 1,336 |
| AMR | 1,482 | 72,284 | 60,295 | 411 |
| EAS | 505 | 27,162 | 12,253 | 116 |
| EUR | 2,072 | 1,371,960 | 330,372 | 12,420 |
| Total | 5,819 | 1,794,714 | 481,186 | 14,283 |

### Optimizations for SAIGE Step One Using GPUs

The primary challenge we encountered when using the SAIGE algorithm on the DOE OLCF Summit HPC was the IBM POWER9 processors incompatibility with the Intel threading building block (TBB) library, which is instrumental for parallelization within step one. This issue prevented us from installing the native SAIGE version, prompting us to find alternative solutions. In addition, solving the logistic mixed model using the PCG algorithm posed challenges due to the numerous iterations and the time-consuming nature of the process. Step one’s time complexity (𝑂) is O(MN^1.5^), where *N* is the sample size, and *M* is the number of genetic markers per sample, making the calculation of the GRM a substantial contributor to the overall computational time for this step, particularly when dealing with large sample sizes (as indicated in equations 1, 2, 3) (10). Building and storing the GRM demanded substantial memory and computational resources. SAIGE’s approach addressed the memory issue by generating GRM segments on-demand, albeit at the cost of increased time requirements and the need for extensive parallelization using multiple CPUs.

SAIGE models the relationship between traits (***Y***) and genotypes (***G***) while adjusting for other covariates (***X)*** and random genetic effects (𝒃) accounting for unknown sample relatedness based on the linear and logistic mixed models (9) (equation 1):

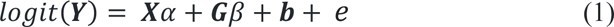

𝛼 and 𝛽 are the coefficient vectors of fixed effects and genetic effects, respectively and *e* is a random effect of residual errors. Each element in 𝒀 represents the probability for an individual being a case given the covariates and genotypes as well as the random effect. The variable ***b*** is assumed to be sampled from a normal distribution with a mean of zero, and a standard deviation of 𝜏𝝍, where 𝝍 is the GRM calculated as

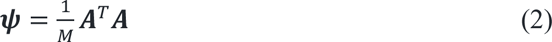

where ***A*** is the genotype matrix of size 𝑁 × 𝑀. Optimizing a linear system solution involving the GRM (𝝍) matrix on GPUs is our focus.

The model is fit under the null hypothesis of 𝛽 = 0, in which the iterative PCG method is used to obtain a solution to a linear system of equations defined by 𝝍𝒙 = 𝒃 for a given vector 𝑏. This iterative process, central to step one, was time-consuming due to multiple matrix-vector operations involving the GRM. Furthermore, building the GRM itself is increasingly memory intensive as the number of individuals and marker panels increase in size. For example, the MVP release 4 population group similar to 1KG-Europeans (*N* = 445,444; *M* = 120,000) would produce a GRM of approximately 800 gigabytes.

To accelerate the computation time and reduce the memory footprint, we employed distributed computing techniques involving the use of Message Passing Interface (MPI) (30) and were able to successfully exploit the parallel computing capability of GPUs for matrix-vector multiplications. Specifically, we partition the columns of the matrix 𝐴 that are used to form the GRM and distribute them into a set of nodes on a cluster. For example, node 𝑖 contains columns 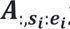 with 𝑠*_i_* and 𝑒*_i_* denoting the start and end indices of the columns of 𝐴 stored in node 𝑖. At each iteration of the PCG method, a matrix-vector multiplication 𝝍𝒗 for some vector 𝒗 is performed. Using the fact that 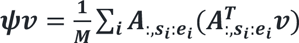, each node computes its summand in parallel on GPUs. The results of all nodes are summed and redistributed using MPI. NVIDIA’s BLAS library *cublasgemv* (30) is used to compute the summand to further accelerate the two matrix-vector multiplications, 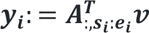 and 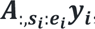 on GPUs (**Figure 2**).

**Fig. 2.**
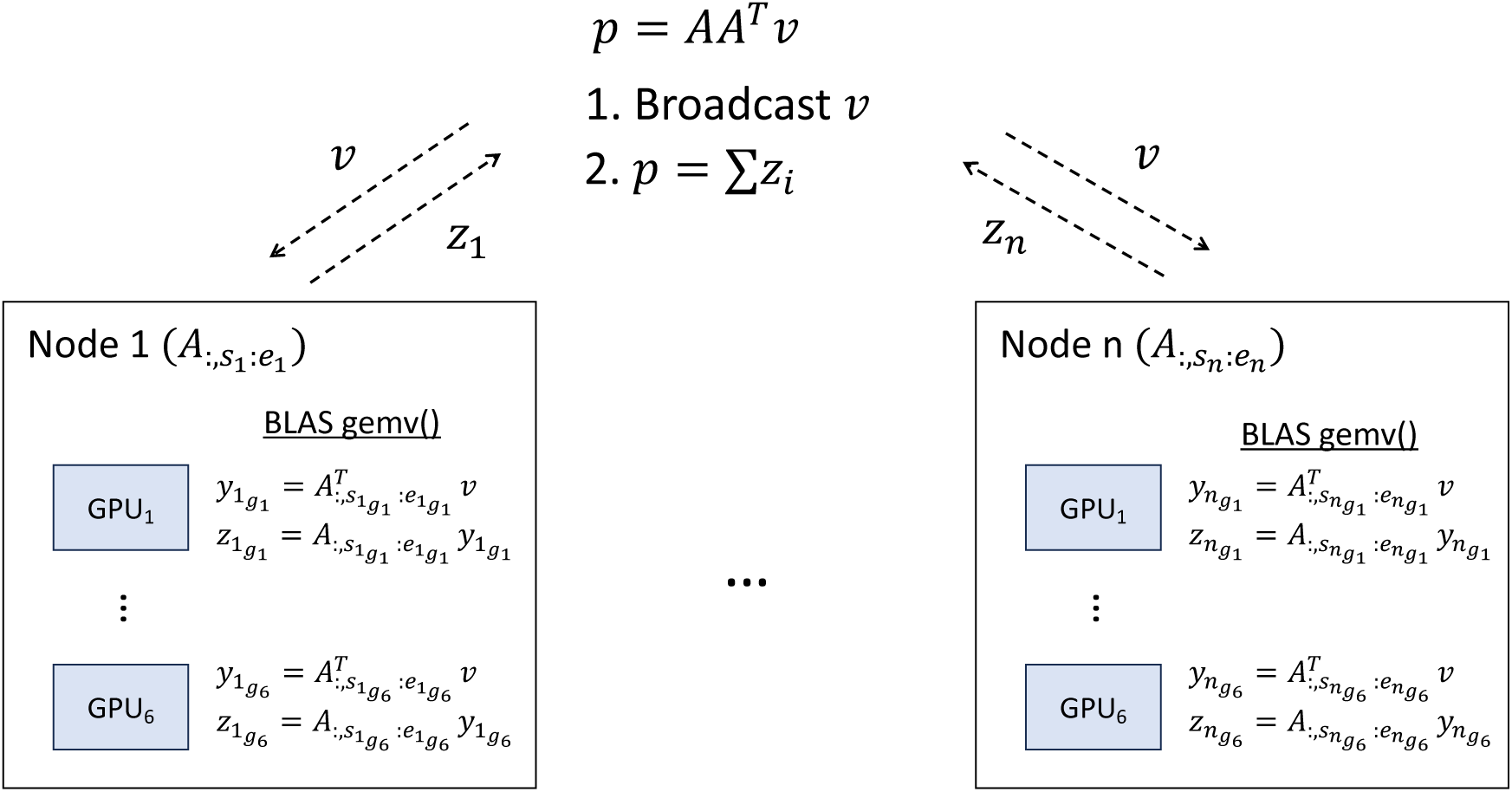
Distributed BLAS gemv(), matrix-vector multiplication, using GPUs on the cluster. The columns of matrix 𝐴 are distributed and preloaded on GPUs, with node 𝑖 having columns with indices from 𝑠_𝑖_ to 𝑒_𝑖_, and these columns are distributed on GPUs on that node. To compute 𝑝 = 𝐴𝐴*^T^*𝑣, we first broadcast 𝑣 to GPUs, and each node computes a partial solution on GPUs. These partial solutions are aggregated to compute a solution 𝑝.

To deal with the large memory requirement, SAIGE relied on the Intel TBB package to parallelize this step, which was incompatible with the Summit infrastructure. We initially replaced the TBB’s parallelization method with OpenMP (32) for executing the matrix-vector operations. However, the primary benefit of accelerating step one lies in the considerably faster matrix computations achieved using GPUs compared to CPUs. We compared the SAIGE version that leveraged OpenMP API for parallelization with the GPU version (**Table 3**) using the Varicose Veins trait (454.1 ICD-9 code). In the OpenMP version, we utilized all 42 available cores on the compute node for parallelizing the matrix calculations to generate the GRM, while for the GPU version we utilized 16 GPUs each equipped with 16 GB of memory in each GPU to distribute subsections of the matrix with dimensions of 8,256 by 445,444. On average, a single PCG iteration on a GPU required approximately 0.069 seconds for the group similar to 1KG-Europe in MVP. In contrast, the OpenMP SAIGE implementation took roughly 5.06 seconds, marking a substantial 72-fold improvement for PCG iterations to converge (**Figure 3**). It took 30 minutes (on 3 nodes with 6 GPUs each) using the GPU-SAIGE implementation to complete step one. Conversely, the same analysis conducted with the OpenMP implementation took 4 hours and 8 minutes in a single 42-core node, representing an overall 3-fold improvement and considers other processes within step one such as processing the input data. While the OpenMP implementation successfully executed the calculations in SAIGE step one on CPUs with a low memory footprint, it required numerous CPU parallel processes to achieve convergence. The advantage of using GPUs becomes readily apparent as the genotype matrix size grows, because it takes substantially longer for CPUs to parallelize the matrix operations. The GPU capitalizes on its inherent parallelization capabilities and pre-loading contents of the matrix into memory, offering a substantial performance boost for large-scale genetic analysis.

**Fig. 3.**
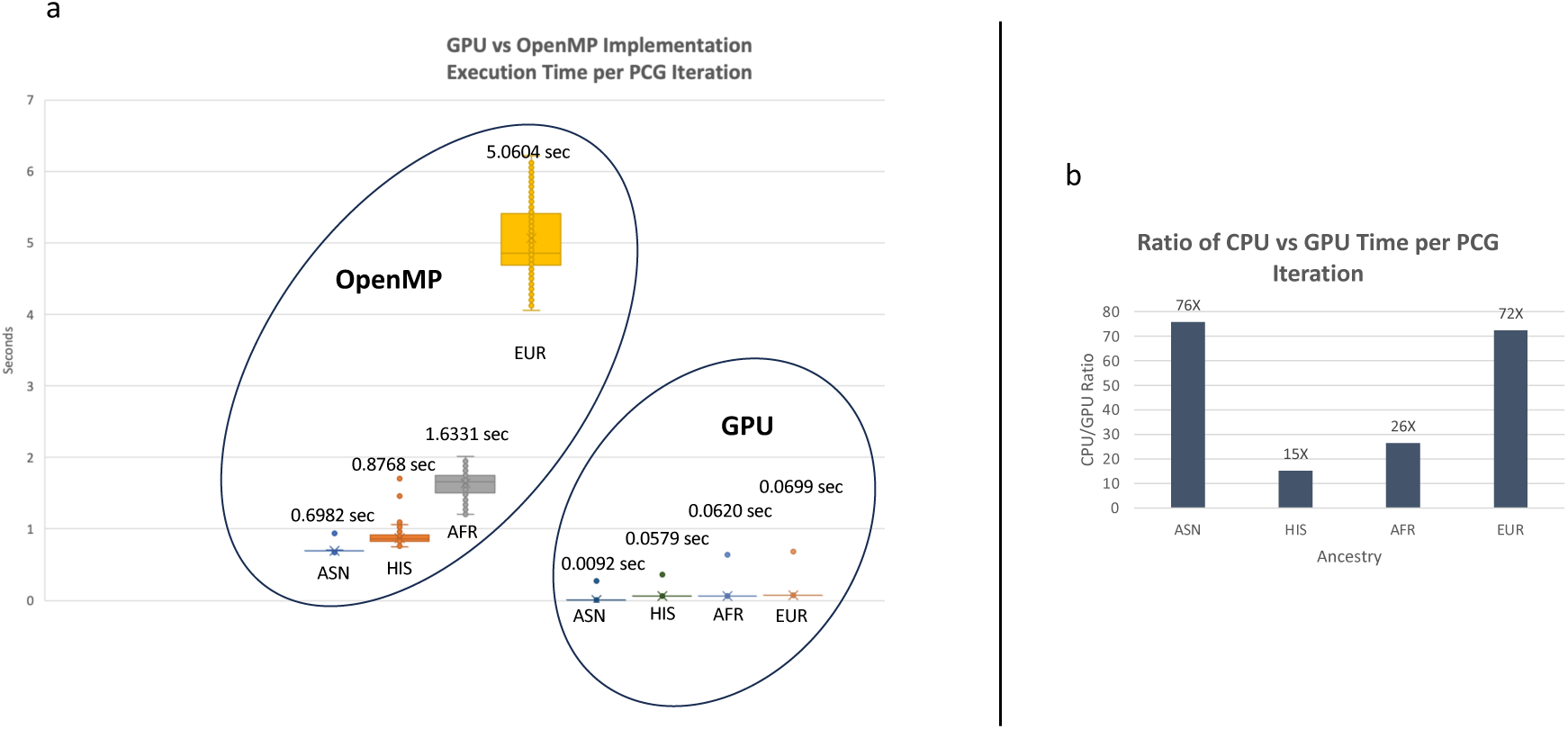
Comparative Performance of GPU and CPU Implementations in SAIGE Step one - This figure compares the execution time for each iteration of matrix operations in SAIGE Step one for the European population group. a) Demonstration of the time required for a single PCG iteration on a GPU, showcasing the efficient parallelization within the GPU. b) Contrast with the OpenMP implementation on CPUs, emphasizing the significant speed improvement achieved with GPU acceleration. As the genotype matrix size increases, the advantage of using the GPU version becomes more pronounced, as highlighted by the diminishing execution time on the GPU compared to the CPU.

**Table 3.** Execution times for SAIGE step one on Varicose Veins (ICD-9 code 454.1) using 3 versions of the SAIGE algorithm on the different OLCF infrastructures. CPU environment contained 32-core nodes, while the GPU nodes contain 42-cores and GPUs with 32 GB of RAM.

| Population Group | Subjects | Step one for Varicose Veins (hours) |  |  |
| --- | --- | --- | --- | --- |
|  |  | Native SAIGE | SAIGE-OpenMP | SAIGE-GPU |
| AFR | 121,725 | 5.10 | 1.06 | 0.38 |
| AMR | 51,124 | 1.50 | 0.97 | 0.28 |
| EAS | 8,003 | 0.97 | 0.58 | 0.23 |
| EUR | 458,307 | 25.75 | 4.10 | 1.50 |

It is important to note that, due to computing the complete GRM in parallel GPUs, the memory footprint increased in comparison to the CPU-based approach which processes the GRM in small segments independent of one another. Thus, the number of nodes needed to cover the GPUs is increased per trait in larger population groups. The amount GPUs required for a run was calculated using the formula:

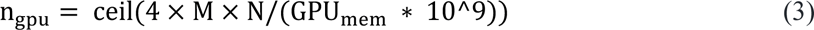

This calculation factored in GPU memory capacity (GPU_454_), the byte size of a single precision floating-point number (4 bytes), and the conversion between bytes and gigabytes (10^9^). This formula can be used for any cohort in additional biobanks to determine the number of GPUs to be used on other computational environments (i.e., cloud infrastructure). This estimation considered the linear relationship between the genotype matrix size, GPU memory available and the required number of nodes which can be visualized (**Figures 4a, 4b**).

**Fig. 4.**
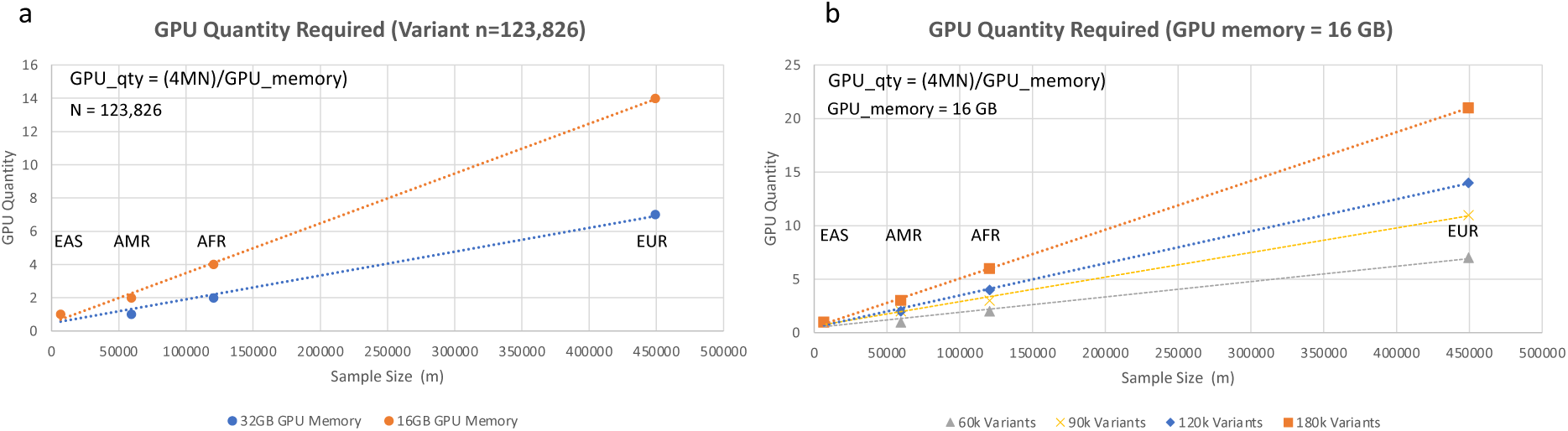
GPU Node Requirements and Memory Impact – GPU node requirement highlight the linear relationship between genotype matrix size and the required number of nodes, offering insights into efficient GPU utilization. The GPU node requirement factored in the GPU memory, the byte size of a single precision floating-point number, and the conversion between bytes and gigabytes. A) Impact of changing the memory available in the GPU. B) Impact of changing number of genotype variants in the input matrix and fixing the GPU memory to 16 gigabytes per GPU, emphasizing considerations for diverse biobank cohorts and computational environments.

The optimizations made in step one effectively harnessed the speed of GPU matrix computation and parallelization, resulting in a significant reduction in analysis time. The GPU optimization of step one enabled the completion of the GWAS analysis for all traits and population groups within 2,381 node hours, representing a remarkable 20-fold improvement for step one in comparison to the initial native SAIGE implementation in a CPU-based cluster (as presented in **Table 2**). Consequently, step one was accomplished in less than 5 days through efficient utilization of node hours facilitated by high-memory Summit nodes for all MVP traits and population groups. Overall, an effective usage of 22,051 GPUs was needed to complete the analysis.

### SAIGE Step two Job Management

In step two of the SAIGE algorithm, millions of variant association tests were conducted independently, given the highly parallel nature of these jobs. Execution times for both the SAIGE-GPU and SAIGE-OpenMP implementations incorporated these optimizations for step two which showed an improvement of 2 to 3-fold compared to initial tests (as summarized in **Table 4)**. To enable parallelization, the MVP genotype data files were partitioned into 219 files based on imputation analysis results. This data partitioning strategy facilitated the parallel execution of 219 jobs per trait and population group, totaling nearly 2 million independent jobs. The predominant challenge in step two revolved around managing the substantial number of jobs required for which we used the R library Tasktools (33), enabling successful submission and monitoring of the jobs.

**Table 4.**
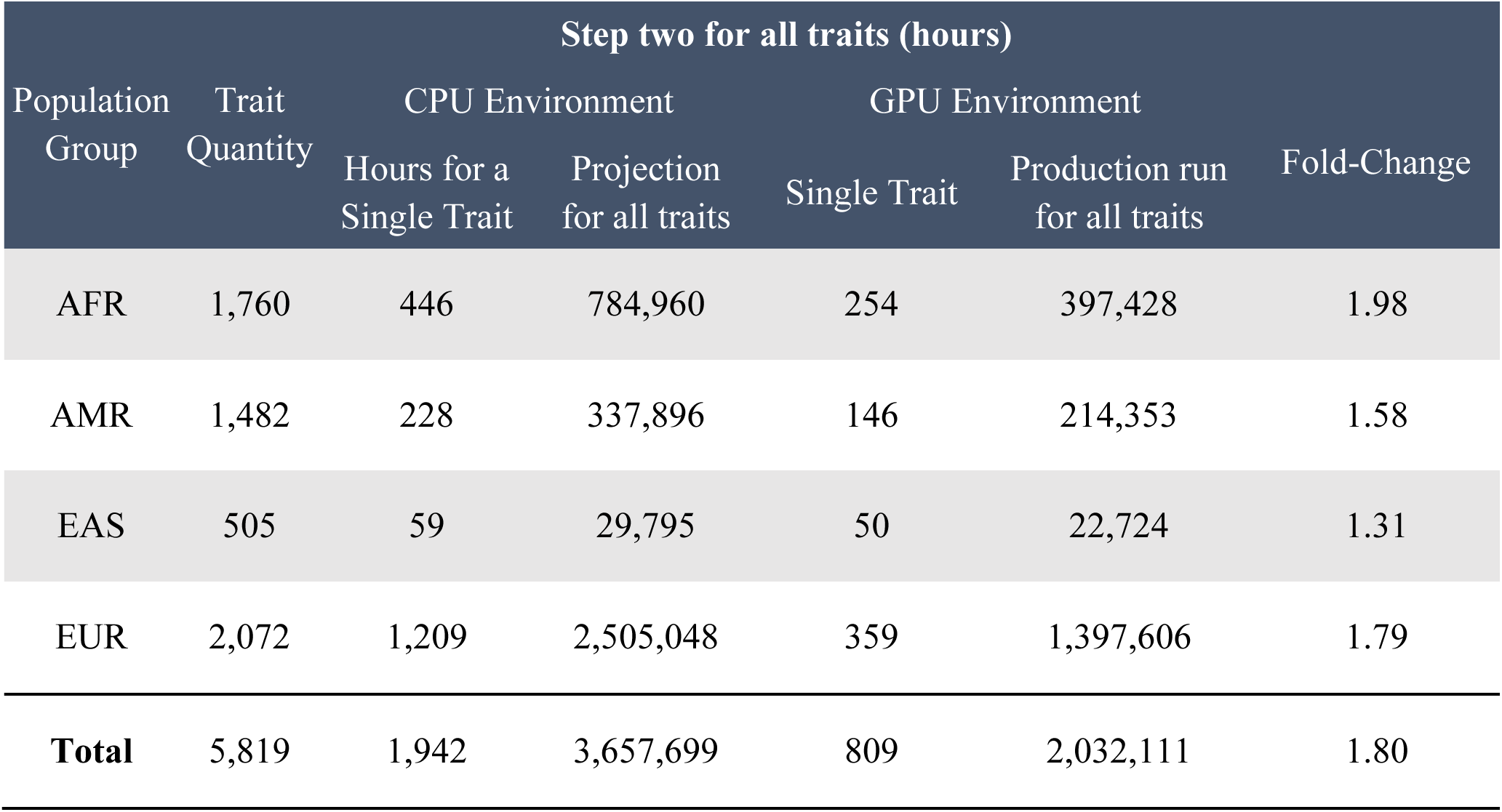
Execution time for SAIGE step two on Varicose Veins (PheCode 454.1) using 2 versions of the SAIGE algorithm on a CPU and GPU environment.

| Step two for all traits (hours) |  |  |  |  |  |  |
| --- | --- | --- | --- | --- | --- | --- |
| Population Group | Trait Quantity | CPU Environment |  | GPU Environment |  | Fold-Change |
|  |  | Hours for a Single Trait | Projection for all traits | Single Trait | Production run for all traits |  |
| AFR | 1,760 | 446 | 784,960 | 254 | 397,428 | 1.98 |
| AMR | 1,482 | 228 | 337,896 | 146 | 214,353 | 1.58 |
| EAS | 505 | 59 | 29,795 | 50 | 22,724 | 1.31 |
| EUR | 2,072 | 1,209 | 2,505,048 | 359 | 1,397,606 | 1.79 |
| **Total** | 5,819 | 1,942 | 3,657,699 | 809 | 2,032,111 | 1.80 |

### SAIGE-GPU Container on Cloud Infrastructures

While the comprehensive analysis was conducted on the OLCF Summit HPC infrastructure, as the MVP cohort data was exclusively available on OLCF computational resources, we note that other cohorts, such as the United Kingdom Biobank (UKBB) (3) and All of Us Biobank (AoU) (7), can only be accessed through cloud infrastructures like the Google Cloud Platform (GCP) and Azure. In response to this demand, we have developed a specialized container image designed for versatile deployment across various cloud infrastructures.

To evaluate its performance, we conducted a comparative study that pitted SAIGE-GPU against SAIGE-CPU using data from the UK and AoU Biobanks. We employed the Type 2 Diabetes (T2D) trait to assess their precision, processing speed, and cost-effectiveness within the GCP cloud environment for two of the largest genetically inferred population groups, namely African and European (**Figure 5** and **Table 5**). For instance, a 5-fold improvement in execution time was seen when analyzing the T2D trait from AoU across the European population group (N = 133,000; M = 100,000). Step one completed in 10 minutes using 1 GPU (A100 GPU, 85 GB RAM), whereas the CPU-based SAIGE version consumed 45 minutes on a 64-core virtual machine. Furthermore, the cost of utilizing 1 GPU for the EUR cohort amounted to approximately $0.42, while the cost of the 64-core VM was $3.17. A similar trend in terms of cost and time is observed for the AFR population group, which would have a smaller memory footprint due to the matrix size.

**Fig. 5.**
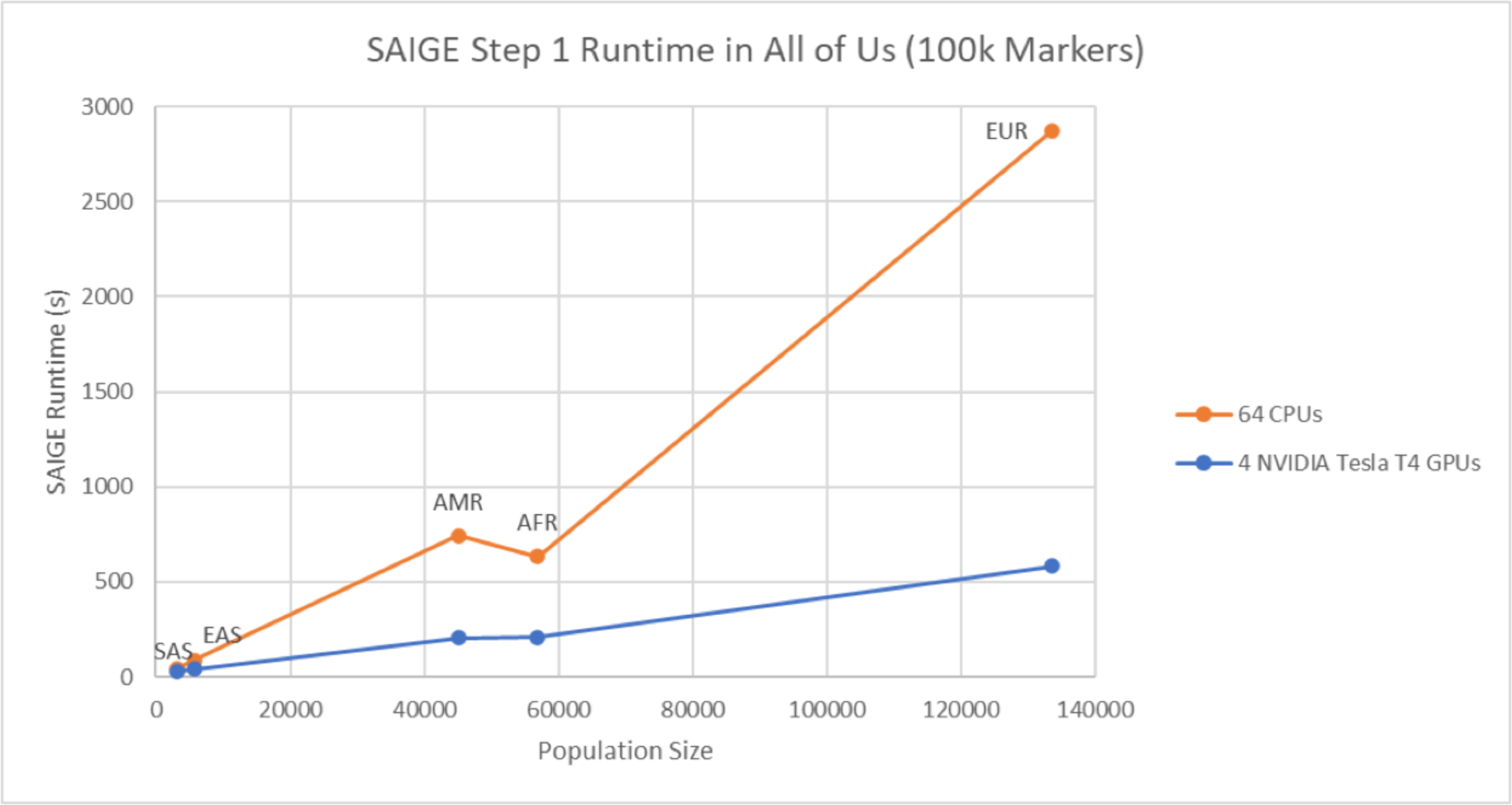
SAIGE step one run time for All of Us data. The figure shows the time comparison of running SAIGE step one for the T2D phenotype on the Google Cloud Platform for the 5 population groups (EUR, AFR, AMR, EAS, SAS). The analysis was executed on 4 NVIDIA T4 GPUs for the SAIGE-GPU version and a 64-CPU VM for the SAIGE-CPU version.

**Table 5.** Cost and time execution comparison using All of Us and UK Biobank data on Google Cloud Platform for SAIGE-GPU version vs the native SAIGE version.

| Category | All of Us |  | UK Biobank |  |
| --- | --- | --- | --- | --- |
|  | European<sup>+</sup> | African American<sup>+</sup> | European<sup>+</sup> | African<sup>+</sup> |
| Variant Size | 100,000 | 100,000 | 100,000 | 100,000 |
| Sample Size | 133,000 | 55,000 | 420,500 | 6,600 |
| SAIGE GPU Analysis Time (hours)* | 0.16 | 0.1 | 0.55 | 0.02 |
| SAIGE GPU Analysis Cost | \$0.42 | \$0.26 | \$1.45 | \$0.05 |
| SAIGE CPU Analysis Time (hours)** | 0.8 | 0.17 | 0.98 | 0.25 |
| SAIGE CPU Analysis Cost | \$3.17 | \$0.67 | \$3.88 | \$0.99 |

\* Google Cloud - A100 GPU, 85 GB RAM, \$2.64/hour
\*\* Google Cloud - 96 Core VM, \$3.96/hour
+ Phenotype used was Type 2 Diabetes

This same pattern of advantages is evident when applied to UKBB traits, as exemplified in table 5. Specifically, we focused on the EUR population group, which consisted of 420,500 individuals, closely resembling the MVP EUR cohort in participant size. GCP infrastructure (NVIDIA Tesla A100 GPUs, 12 vCPUs, and 85GB of RAM) was employed to run the T2D trait and completed the analysis in just over 30 minutes, with an average cost of $1.45. In contrast, utilizing the CPU-based SAIGE version consumed 58 minutes and incurred a cost of $3.88 using a 96-core VM.

### Conclusion

We leveraged the GPU computational resources of the DOE OLCF Summit HPC address major computational challenges posed by the increasingly large datasets utilized in genomics research. In this example, we demonstrate optimization of a highly used tool for genomic analyses designed for CPUs, SAIGE. Prior to optimizing step one with GPUs, the analysis would have spanned several years for all genetically inferred population groups. The optimizations have now condensed the completion time to under a month, reducing node hours by a substantial factor.

Even though we largely focused on step one of SAIGE, in Step two we showed how we executed millions of variant association tests in parallel, a highly compute-intensive task. We intend to further improve this step by parallelizing this step instead of performing each association in serial mode.

In a recent article (34), the authors performed a large analysis on over 7,000 traits of the Pan-UK Biobank (35) data for multiple ancestries using hail-batch on the Google Cloud Platform. As previously mentioned, both the European population groups for UKBB and MVP are comparable in size, while the African, Admixed American and Eastern Asian population groups are larger for the MVP. The authors used the SAIGE-CPU implementation to perform close to 300 billion associations and required over 3.8 million CPU hours to complete both step one and step two in SAIGE. In comparison, the MVP analysis required 14,283 GPU hours for step one and approximately 2 million CPU hours for step two to perform over 350 billion associations.

The MVP has now expanded to a million individuals (36) and plans to collect whole-genome sequencing data, likely to increase the number of low-frequency variants that will be tested in the future. Thus, it is imperative to understand approaches to efficiently optimize software already developed for these data in HPC environments. Our primary focus lay in enhancing the efficiency of SAIGE’s first step since it is iteratively employed in numerous downstream SAIGE-related analyses (e.g., SAIGE-GENE (37)). However, our ongoing efforts center on further streamlining SAIGE for GW-PheWAS studies across multiple biobanks such as All of Us, UK Biobank, Penn Medicine BioBank (2).

The continuous evolution of GPU technology in various implementations offers a promising outlook. The Summit infrastructure currently harnesses NVIDIA CUDA libraries for these operations, but future systems may incorporate different libraries, further accelerating execution times and lowering costs. These systems are expected to feature expanded memory and storage capacities. Additionally, our GPU-based SAIGE implementation can be readily adapted for Intel GPUs using the Intel oneAPI platform and AMD GPUs using their ROCm platform.

A container is available for deployment on cloud platforms equipped with GPU nodes. The code can be accessed at https://exascale-genomics.github.io/SAIGE-GPU. The significant improvements in efficiency achieved with SAIGE using GPUs demonstrate the potential for the development of new and existing tools capable of performing population analysis at the exascale level by optimizing software for GPU usage.

## Supporting information

MVP_Author_list_supplemental

## Acknowledgments

We thank the Million Veteran Program, Office of Research and Development, and Veterans Health Administration for supporting this work. We would like to sincerely thank Dr. Thomas Zacharia for providing access to the supercomputers at the Oak Ridge National Laboratory Leadership Computing Facility and Dr. Dimitri Kusenov, the previous DOE Headquarters lead for the VA-DOE partnership, for his invaluable guidance and support. Their contributions have been instrumental in the successful completion of this study. Last but not least, we thank former staff members, and volunteers, who have contributed to MVP and, most of all, MVP participants for their service and their continued contributions to our nation through participation in this study. This publication does not represent the views of the Department of Veteran Affairs or the United States Government.

## Author contributions

**Conceptualization**: AAR, YK, TNN, KK, RB, JEH, MC, ML, KM, JH, DS, EB, GT, SM, KC, MJG, BFV, SD, KPL, WZ, AV, RKM; **Methodology**: AAR, YK, TNN, KK, RB, JEH, MC, ML, KM, JH, DS, BFV, SD, KPL, WZ, AV, RKM; **Investigation**: AAR, YK, TNN, KK, RB, JEH, MC, ML, KM, JH, DS, MC, SM, BFV, KC, MJG, SD, KPL, WZ, AV,RKM; **Visualization**: AAR, YK, TNN, KK, JEH, MC, SM, BFV, KC, MJG, SD, KPL, WZ, AV, RKM**; Funding acquisition:** EB, GT, PN, SM, KC, MJG, SD, KPL, AV, RKM; **Project administration:** AAR, JEH, MJG, SD, KPL, WZ, AV, RKM; **Supervision:** EB, GT, SM, PN, AV, MJG, SD, KPL, RKM; **Writing – original draft:** AAR, YK, TNN, KK, RB, JEH, AV, WZ, RKM; **Writing – review & editing:** AAR, YK, TNN, KK, RB, JEH, MC, ML, SM, BFV, KC, MJG, SD, KPL, WZ, AV, RKM.

## Data and materials availability

The optimized SAIGE-GPU software can be accessed at the GitHub page https://exascale-genomics.github.io/SAIGE-GPU.

