## Supplementary material for "Accelerating Genome- and Phenome-Wide Association Studies using GPUs – A case study using data from the Million Veteran Program": MVP_Author_list_supplemental

**VA Million Veteran Program:**

**Core Acknowledgement for Publications**

**February 2024**

**MVP Program Office**

- Sumitra Muralidhar, Ph.D., Program Director

US Department of Veterans Affairs, 810 Vermont Avenue NW, Washington, DC 20420

- Jennifer Moser, Ph.D., Associate Director, Scientific Programs

US Department of Veterans Affairs, 810 Vermont Avenue NW, Washington, DC 20420

- Jennifer E. Deen, B.S., Associate Director, Cohort & Public Relations

US Department of Veterans Affairs, 810 Vermont Avenue NW, Washington, DC 20420

**MVP Executive Committee**

- Co-Chair: Philip S. Tsao, Ph.D.

VA Palo Alto Health Care System, 3801 Miranda Avenue, Palo Alto, CA 94304

- Co-Chair: Sumitra Muralidhar, Ph.D.

US Department of Veterans Affairs, 810 Vermont Avenue NW, Washington, DC 20420

- J. Michael Gaziano, M.D., M.P.H.

VA Boston Healthcare System, 150 S. Huntington Avenue, Boston, MA 02130

- Elizabeth Hauser, Ph.D.

Durham VA Medical Center, 508 Fulton Street, Durham, NC 27705

- Amy Kilbourne, Ph.D., M.P.H.

VA HSR&D, 2215 Fuller Road, Ann Arbor, MI 48105

- Michael Matheny, M.D., M.S., M.P.H.

VA Tennessee Valley Healthcare System, 1310 24^th^ Ave. South, Nashville, TN 37212

- Dave Oslin, M.D.

Philadelphia VA Medical Center, 3900 Woodland Avenue, Philadelphia, PA 19104

**MVP Co-Principal Investigators**

- J. Michael Gaziano, M.D., M.P.H.

VA Boston Healthcare System, 150 S. Huntington Avenue, Boston, MA 02130

- Philip S. Tsao, Ph.D.

VA Palo Alto Health Care System, 3801 Miranda Avenue, Palo Alto, CA 94304

**MVP Core Operations**

- Jessica V. Brewer, M.P.H., Director, Cohort Operations

VA Boston Healthcare System, 150 S. Huntington Avenue, Boston, MA 02130

- Mary T. Brophy M.D., M.P.H., Director, Biorepository

VA Boston Healthcare System, 150 S. Huntington Avenue, Boston, MA 02130

- Kelly Cho, M.P.H, Ph.D., Director, MVP Phenomics

VA Boston Healthcare System, 150 S. Huntington Avenue, Boston, MA 02130

- Lori Churby, B.S., Director, Regulatory Affairs

VA Palo Alto Health Care System, 3801 Miranda Avenue, Palo Alto, CA 94304

- Scott L. DuVall, Ph.D., Director, VA Informatics and Computing Infrastructure (VINCI)

VA Salt Lake City Health Care System, 500 Foothill Drive, Salt Lake City, UT 84148

- Saiju Pyarajan Ph.D., Director, Data and Computational Sciences

VA Boston Healthcare System, 150 S. Huntington Avenue, Boston, MA 02130

- Luis E. Selva, Ph.D., Director, MVP Biorepository Coordination

VA Boston Healthcare System, 150 S. Huntington Avenue, Boston, MA 02130

- Shahpoor (Alex) Shayan, M.S., Director, MVP PRE Informatics

VA Boston Healthcare System, 150 S. Huntington Avenue, Boston, MA 02130

- Stacey B. Whitbourne, Ph.D., Director, MVP Cohort Development and Management

VA Boston Healthcare System, 150 S. Huntington Avenue, Boston, MA 02130

- MVP Coordinating Centers
  - MVP Coordinating Center, Boston - J. Michael Gaziano, M.D., M.P.H.

VA Boston Healthcare System, 150 S. Huntington Avenue, Boston, MA 02130

- MVP Coordinating Center, Palo Alto – Philip S. Tsao, Ph.D.

VA Palo Alto Health Care System, 3801 Miranda Avenue, Palo Alto, CA 94304

- MVP Information Center, Canandaigua – Brady Stephens, M.S.

Canandaigua VA Medical Center, 400 Fort Hill Avenue, Canandaigua, NY 14424

- Cooperative Studies Program Clinical Research Pharmacy Coordinating Center, Albuquerque – Todd Connor, Pharm.D.; Dean P. Argyres, B.S., M.S.

New Mexico VA Health Care System, 1501 San Pedro Drive SE, Albuquerque, NM 87108

**MVP Publications and Presentations Committee**

- Co-Chair: Themistocles L. Assimes, M.D., Ph. D

VA Palo Alto Health Care System, 3801 Miranda Avenue, Palo Alto, CA 94304

- Co-Chair: Adriana Hung, M.D.; M.P.H

VA Tennessee Valley Healthcare System, 1310 24^th^ Ave. South, Nashville, TN 37212

- Co-Chair: Henry Kranzler, M.D.

Philadelphia VA Medical Center, 3900 Woodland Avenue, Philadelphia, PA 19104

**MVP Local Site Investigators**

- Samuel Aguayo, M.D., Phoenix VA Health Care System

650 E. Indian School Road, Phoenix, AZ 85012

- Sunil Ahuja, M.D., South Texas Veterans Health Care System

7400 Merton Minter Boulevard, San Antonio, TX 78229

- Kathrina Alexander, M.D., Veterans Health Care System of the Ozarks

1100 North College Avenue, Fayetteville, AR 72703

- Xiao M. Androulakis, M.D., Columbia VA Health Care System

6439 Garners Ferry Road, Columbia, SC 29209

- Prakash Balasubramanian, M.D., William S. Middleton Memorial Veterans Hospital

2500 Overlook Terrace, Madison, WI 53705

- Zuhair Ballas, M.D., Iowa City VA Health Care System

601 Highway 6 West, Iowa City, IA 52246-2208

- Elizabeth S. Bast, M.D., M.P.H., Miami VA Health Care System

1201 NW 16th Street, 11 GRC, Miami FL 33125

- Jean Beckham, Ph.D., Durham VA Medical Center

508 Fulton Street, Durham, NC 27705

- Sujata Bhushan, M.D., VA North Texas Health Care System

4500 S. Lancaster Road, Dallas, TX 75216

- Edward Boyko, M.D., VA Puget Sound Health Care System

1660 S. Columbian Way, Seattle, WA 98108-1597

- David Cohen, M.D., Portland VA Medical Center

3710 SW U.S. Veterans Hospital Road, Portland, OR 97239

- Louis Dellitalia, M.D., Birmingham VA Medical Center

700 S. 19th Street, Birmingham AL 35233

- Gerald Wayne Dryden, Jr., M.D., Ph.D., Louisville VA Medical Center

800 Zorn Avenue, Louisville, KY 40206

- L. Christine Faulk, M.D., Robert J. Dole VA Medical Center

5500 East Kellogg Drive, Wichita, KS 67218-1607

- Joseph Fayad, M.D., VA Southern Nevada Healthcare System

6900 North Pecos Road, North Las Vegas, NV 89086

- Daryl Fujii, Ph.D., VA Pacific Islands Health Care System

459 Patterson Rd, Honolulu, HI 96819

- Saib Gappy, M.D., John D. Dingell VA Medical Center

4646 John R Street, Detroit, MI 48201

- Frank Gesek, Ph.D., White River Junction VA Medical Center

163 Veterans Drive, White River Junction, VT 05009

- Michael Godschalk, M.D., Richmond VA Medical Center

1201 Broad Rock Blvd., Richmond, VA 23249

- Jennifer Greco, M.D., Sioux Falls VA Health Care System

2501 W 22nd Street, Sioux Falls, SD 57105

- Todd W. Gress, M.D., Ph.D., Hershel “Woody” Williams VA Medical Center

1540 Spring Valley Drive, Huntington, WV 25704

- Samir Gupta, M.D., M.S.C.S., VA San Diego Healthcare System

3350 La Jolla Village Drive, San Diego, CA 92161

- Salvador Gutierrez, M.D., Edward Hines, Jr. VA Medical Center

5000 South 5th Avenue, Hines, IL 60141

- Mark Hamner, M.D., Ralph H. Johnson VA Medical Center

109 Bee Street, Mental Health Research, Charleston, SC 29401

- John Harley, M.D., Ph.D., Cincinnati VA Medical Center

3200 Vine Street, Cincinnati, OH 45220

- Daniel J. Hogan, M.D., Bay Pines VA Healthcare System

10,000 Bay Pines Blvd Bay Pines, FL 33744

- Adriana Hung, M.D., M.P.H., VA Tennessee Valley Healthcare System

1310 24th Avenue, South Nashville, TN 37212

- Robin Hurley, M.D., W.G. (Bill) Hefner VA Medical Center

1601 Brenner Ave, Salisbury, NC 28144

- Pran Iruvanti, D.O., Ph.D., Hampton VA Medical Center

100 Emancipation Drive, Hampton, VA 23667

- Frank Jacono, M.D., VA Northeast Ohio Healthcare System

10701 East Boulevard, Cleveland, OH 44106

- Darshana Jhala, M.D., Philadelphia VA Medical Center

3900 Woodland Avenue, Philadelphia, PA 19104

- Seema Joshi, M.D., F.A.C.P., ABOIM; VA Eastern Kansas Health Care System

4101 S 4th Street Trafficway, Leavenworth, KS 66048

- Scott Kinlay, M.B.B.S., Ph.D., VA Boston Healthcare System

150 S. Huntington Avenue, Boston, MA 02130

- Michael Landry, Ph.D., Southeast Louisiana Veterans Health Care System

2400 Canal Street, New Orleans, LA 70119

- Peter Liang, M.D., M.P.H., VA New York Harbor Healthcare System

423 East 23rd Street, New York, NY 10010

- Suthat Liangpunsakul, M.D., M.P.H., Richard Roudebush VA Medical Center

1481 West 10th Street, Indianapolis, IN 46202

- Jack Lichy, M.D., Ph.D., Washington DC VA Medical Center

50 Irving St, Washington, D. C. 20422

- Tze Shien Lo, M.D., Fargo VA Health Care System

2101 N. Elm, Fargo, ND 58102

- C. Scott Mahan, M.D., Charles George VA Medical Center

1100 Tunnel Road, Asheville, NC 28805

- Ronnie Marrache, M.D., VA Maine Healthcare System Center, Augusta, ME 04330
- Stephen Mastorides, M.D., James A. Haley Veterans’ Hospital

13000 Bruce B. Downs Blvd, Tampa, FL 33612

- Kristin Mattocks, Ph.D., M.P.H., Central Western Massachusetts Healthcare System

421 North Main Street, Leeds, MA 01053

- Paul Meyer, M.D., Ph.D., Southern Arizona VA Health Care System

3601 S 6th Avenue, Tucson, AZ 85723

- Jonathan Moorman, M.D., Ph.D., James H. Quillen VA Medical Center

Corner of Lamont & Veterans Way, Mountain Home, TN 37684

- Providencia Morales, R.N., Northern Arizona VA Health Care System

500 Highway 89 North, Prescott, AZ 86313

- Timothy Morgan, M.D., VA Long Beach Healthcare System

5901 East 7th Street Long Beach, CA 90822

- Maureen Murdoch, M.D., M.P.H., Minneapolis VA Health Care System

One Veterans Drive, Minneapolis, MN 55417

- Eknath Naik, M.D., Ph.D., West Palm Beach VA Medical Center,

7305 North Military Trail, West Palm Beach, FL 33410-6400

- James Norton, Ph.D., VA Health Care Upstate New York

113 Holland Avenue, Albany, NY 12208

- Olaoluwa Okusaga, M.D., Michael E. DeBakey VA Medical Center

2002 Holcombe Blvd, Houston, TX 77030

- Michael K. Ong, M.D., VA Greater Los Angeles Health Care System

11301 Wilshire Blvd, Los Angeles, CA 90073

- Kris Ann Oursler, M.D., Salem VA Medical Center

1970 Roanoke Blvd, Salem, VA 24153

- Ismene Petrakis, M.D., VA Connecticut Healthcare System

950 Campbell Avenue, West Haven, CT 06516

- Samuel Poon, M.D., Manchester VA Medical Center

718 Smyth Road, Manchester, NH 03104

- Amneet S. Rai, Pharm.D., VA Sierra Nevada Health Care System

975 Kirman Avenue, Reno, NV 89502

- Michael Rauchman, M.D., St. Louis VA Health Care System

915 North Grand Blvd, St. Louis, MO 63106

- Richard Servatius, Ph.D., Syracuse VA Medical Center

800 Irving Avenue, Syracuse, NY 13210

- Satish Sharma, M.D., Providence VA Medical Center

830 Chalkstone Avenue, Providence, RI 02908

- River Smith, Ph.D., Eastern Oklahoma VA Health Care System

1011 Honor Heights Drive, Muskogee, OK 74401

- Peruvemba Sriram, M.D., N. FL/S. GA Veterans Health System

1601 SW Archer Road, Gainesville, FL 32608

- Patrick Strollo, Jr., M.D., VA Pittsburgh Health Care System

University Drive, Pittsburgh, PA 15240

- Neeraj Tandon, M.D., Overton Brooks VA Medical Center

510 East Stoner Ave, Shreveport, LA 71101

- Philip Tsao, Ph.D., VA Palo Alto Health Care System

3801 Miranda Avenue, Palo Alto, CA 94304-1290

- Gerardo Villareal, M.D., New Mexico VA Health Care System

1501 San Pedro Drive, S.E. Albuquerque, NM 87108

- Jessica Walsh, M.D., VA Salt Lake City Health Care System

500 Foothill Drive, Salt Lake City, UT 84148

- John Wells, Ph.D., Edith Nourse Rogers Memorial Veterans Hospital

200 Springs Road, Bedford, MA 01730

- Jeffrey Whittle, M.D., M.P.H., Clement J. Zablocki VA Medical Center

5000 West National Avenue, Milwaukee, WI 53295

- Mary Whooley, M.D., San Francisco VA Health Care System

4150 Clement Street, San Francisco, CA 94121

- Peter Wilson, M.D., Atlanta VA Medical Center

1670 Clairmont Road, Decatur, GA 30033

- Junzhe Xu, M.D., VA Western New York Healthcare System

3495 Bailey Avenue, Buffalo, NY 14215-1199

- Shing Shing Yeh, Ph.D., M.D., Northport VA Medical Center

79 Middleville Road, Northport, NY 11768

- Andrew W. Yen, M.D., VA Northern California Health Care System

10535 Hospital Way, Mather, CA 95655
